# Lack of evidence of ACE2 expression and replicative infection by SARS-CoV-2 in human endothelial cells

**DOI:** 10.1101/2020.12.02.391664

**Authors:** Ian McCracken, Gaye Saginc, Liqun He, Alik Huseynov, Alison Daniels, Sarah Fletcher, Claire Peghaire, Viktoria Kalna, Maarja Andaloussi-Mäe, Lars Muhl, Nicky M. Craig, Samantha J. Griffiths, Jürgen G. Haas, Christine Tait-Burkard, Urban Lendahl, Graeme M. Birdsey, Christer Betsholtz, Michela Noseda, Andrew Baker, Anna M. Randi

**Author notes:** shared first Authorship (Ian McCracken, Gaye Saginc, Liqun He, Alik Huseynov). contributed equally (Alison Daniels, Sarah Fletcher). shared senior authorship (Graeme M. Birdsey, Christer Betsholtz, Michela Noseda, Andrew Baker, Anna M. Randi).

## Abstract

A striking feature of severe COVID-19 is thrombosis in large as well as small vessels of multiple organs. This has led to the assumption that SARS-CoV-2 virus directly infects and damages the vascular endothelium. However, endothelial expression of ACE2, the cellular receptor for SARS-CoV-2, has not been convincingly demonstrated. Interrogating human bulk and single-cell transcriptomic data, we found *ACE2* expression in endothelial cells to be extremely low or absent *in vivo* and not upregulated by exposure to inflammatory agents *in vitro*. Also, the endothelial chromatin landscape at the *ACE2* locus showed presence of repressive and absence of activation marks, suggesting that the gene is inactive in endothelial cells. Finally, we failed to achieve infection and replication of SARS-CoV-2 in cultured human endothelial cells, which were permissive to productive infection by coronavirus 229E that uses CD13 as the receptor. Our data suggest that SARS-Cov-2 is unlikely to infect endothelial cells directly; these findings are consistent with a scenario where endothelial injury is indirectly caused by the infection of neighbouring epithelial cells and/or due to systemic effects mediated by immune cells, platelets, complement activation, and/or proinflammatory cytokines.

## Introduction

A striking feature of severe forms of coronavirus disease 2019 (COVID-19), the current pandemic caused by the coronavirus SARS-CoV-2, is severe endothelial injury with micro- and macro-thrombotic disease in the lung and other organs, including the heart. This has led to speculation that viral infection may damage the endothelium through two mechanisms: indirectly, via neighbourhood effects, circulating mediators and immune mechanisms, or directly by viral infection of endothelial cells (EC).

To support the hypothesis of direct viral damage of EC via virus-induced infection, the cells should express the main receptor for SARS-CoV-2, angiotensin-converting enzyme 2 (ACE2), a metalloproteinase component of the renin-angiotensin hormone system and a critical regulator of cardiovascular homeostasis^1^. Indeed, several recent review articles propose that SARS-CoV-2 binding to ACE2 on EC is the mechanism through which the virus may cause direct endothelial damage and endothelialitis^1^. However, expression of ACE2 in EC has not been convincingly demonstrated to support this assumption, nor has there been sufficient evidence to support a direct infection of EC by SARS-CoV-2.

## Results and Discussion

To address the questions of ACE2 expression in human EC and of the ability of SARS-CoV-2 to infect the endothelium, we interrogated transcriptomic and epigenomic data on human EC and studied the interaction and replication of SARS-Cov-2 and its viral proteins with EC *in vitro*. Analysis of RNA-seq was carried out on ENCODE# data from EC from arterial, venous and microvascular beds, in comparison with epithelial cells from respiratory, gastrointestinal and skin sources. Very low or no basal *ACE2* expression was found in EC, compared to epithelial cells (Figure A-B). Moreover, *in vitro* exposure of EC to inflammatory cytokines reported as elevated in the plasma of patients with severe COVID-19 failed to upregulate ACE2 expression (Figure C).

**Figure:**
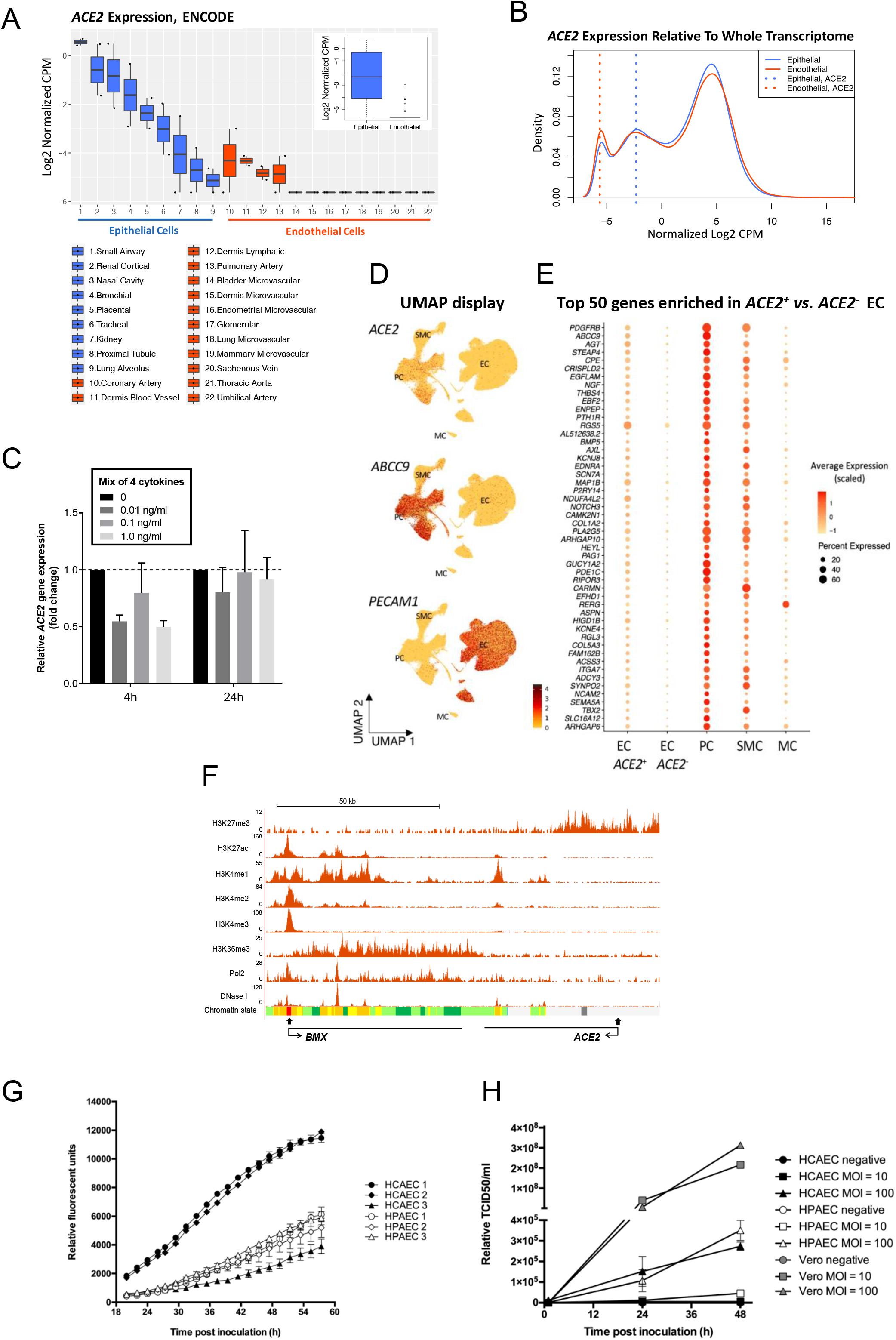
Analysis of *ACE2* expression in human endothelial cells and of coronavirus replication in primary human endothelial cells. **(A-B) Comparison of *ACE2* expression in human primary epithelial and endothelial cells using total RNA-seq data from the ENCODE Database shows low or absent expression in EC.** (A) The difference of *ACE2* expression in epithelial and in endothelial cells is shown in boxplots with individual, as well as grouped samples (inner boxplot). Each dot represents a single sample (n=2 per cell type). (B) Transcriptome profiles of epithelial and endothelial cells are shown in a density plot, using median of all samples per group (n=19360 genes). *ACE2* expression in each group is marked with a dotted line: *ACE2* expression in endothelial cells (red) overlaps with the peak for non-expressed transcripts, while *ACE2* expression in epithelial cells (blue) is to the right, indicating detectable expression. Expression values in all plots are represented in log2 transformed counts per million normalized by Trimmed Mean of M-value (Log2 CPM, blue: epithelial, red: endothelial). **(C) *ACE2* expression is not regulated by inflammatory cytokines in HUVEC.** qPCR analysis of *ACE2* mRNA expression in HUVEC treated with a mix of 4 cytokines/chemokines (TNF-α, IL·l-ß, IL8 and IL6/IL6R chimeric protein) for 4h or 24h at 0, 0.01, 0.1 or 1.0 ng/ml. Data are normalized to *GAPDH* and presented as mean ± SEM of 3 independent experiments. **(D-E) Very low-level, rare and likely contaminating *ACE2* transcripts are seen in EC.** (D) *ACE2* transcript reads are detected preferentially in PC. UMAP landscapes of human heart (ref 2) including 100,579 endothelial cells (EC), 77,856 pericytes (PC), 16,242 smooth muscle cells (SMC) and 718 mesothelial cells (MC). *ACE2* transcript reads are detected preferentially in the PC cluster (enriching for *ABCC9)* and are rare in the EC cluster (enriching for *PECAM1*). (E) PC transcripts are enriched together with *ACE2* in 0.47% of EC. Dot plot displaying the abundance of top-50 transcripts enriched *ACE2^+^* vs. *ACE2*^-^ EC, across cell types indicated in D. (The Wilcoxon Rank Sum tests with Bonferroni-corrected p values are < 1E-60 for each). **(F) Epigenetic profiling indicates that the *ACE2* gene is inactive in EC**. ChIP-seq binding profiles in HUVEC for histone modifications, RNA Pol2 enrichment and DNAse I hypersensitivity. The x axis represents the genomic position, the transcription start sites are indicated by closed arrows and the direction of transcription is indicated by open arrows; the y axis shows ChIP-seq signal in reads per million per base pair (rpm/bp). Chromatin state segmentation colour key: active promoter, red; enhancers, yellow; transcriptional elongation, green; repressed, grey. **(G-H) Coronavirus replication in primary human cardiac and pulmonary endothelial cells shows no replication of SARS-CoV-2**. (G) Viral replication curves in human pulmonary (HPAEC) and cardiac (HCAEC) endothelial cells following infection with HCoV-229E GFP reporter virus (MOI = 0.6). Virus replication was measured via GFP fluorescence every 2 hours from 20 to 58 hours post inoculation. Mean ±SEM of 3 technical replicates are shown at each time point for each biological replicate. (H) Viral growth curves in HPAEC (n=3), HCAEC (n=3), and Vero cells (n=1) following infection with SARS-CoV-2 at MOI = 10 or 100. Supernatant were collected at 1, 24, and 48 hours post infection and virus copy number quantified by RT-qPCR detection of the SARS-CoV-2 N3 gene. TCID= tissue culture infectivity dose.

Single-cell RNA-sequencing (scRNAseq) of human organ donor hearts^2^ showed that while *ACE2* sequence reads are abundant in pericytes (PC), they are rare in EC (Figure D). Out of 100,579 EC, only 468 (0,47%) were *ACE2*^+^, and in the majority (424) only a single *ACE2* transcript was detected. This could reflect true low and rare endothelial *ACE2* expression, but also contamination from adherent PC fragments, a common confounder in vascular scRNAseq data^3^. If such fragments contributed the *ACE2* transcripts observed in certain EC, we would expect to detect other pericyte transcripts in the same cells. Indeed, among the top-50 gene transcripts enriched in *ACE2*^+^ vs. *ACE2*^-^ EC, we noticed several known pericyte markers, including *PDGFRB, ABCC9, KCNJ8* and *RGS5* (Figure E). Comparison of transcript abundance across the three major vascular and mesothelial cells showed that the top-50 gene transcripts were expressed at the highest levels in PC (Figure E). This suggests that the rare occurrence of *ACE2* transcripts in human heart EC is likely caused by pericyte contamination. Similar conclusions have previously been reached in mouse tissues^3^.

Analysis of the chromatin landscape at the *ACE2* gene locus in human umbilical vein EC (HUVEC) using data from ENCODE further supports this concept. The histone modification mark H3K27me3, which indicates repressed chromatin, was enriched at the *ACE2* transcription start site (TSS); conversely, promoter, enhancer and gene body activation marks (H3K27ac, H3K4me1, H3K4me2, H3K4me3, H3K36me3), RNA polymerase-II and DNase I hypersensitivity were absent or low, suggesting that *ACE2* is inactive in EC. In marked contrast, the adjacent gene *BMX*, an endothelial-restricted non-receptor tyrosine kinase displays an epigenetic profile consistent with active endothelial expression (Figure F). Thus, transcriptomic and epigenomic data indicate that *ACE2* is not expressed in human EC.

Other cell surface molecules have been suggested as possible receptors for the virus, but their role in supporting SARS-CoV-2 cell infection remains to be demonstrated. We therefore tested directly whether EC could be capable of supporting coronavirus replication *in vitro*. Productive levels of replication in primary human cardiac and pulmonary EC were observed for the human coronavirus 229E GFP reporter virus^4^, which utilises CD13 as its receptor, demonstrating directly that human EC can support coronavirus replication in principle (Figure G). However, when cells were exposed to SARS-CoV-2, replication levels were extremely low for EC, even following exposure to very high concentrations of virus compared to more permissive VeroE6 cells (Figure H). The observed low levels of SARS-CoV-2 replication in EC are likely explained by viral entry via a non-ACE2 dependent route, due to exposure to supraphysiological concentrations of virus in these experiments (MOI 10 and 100).

These data indicate that direct endothelial infection by SARS-Cov-2 is not likely to occur. The endothelial damage reported in severely ill COVID19 patients is more likely secondary to infection of neighbouring cells and/or other mechanisms, including immune cells, platelets and complement activation, and circulating proinflammatory cytokines. Our hypothesis is corroborated by recent evidence that plasma from critically ill and convalescent patients with COVID-19 causes endothelial cell cytotoxicity^5^. These finding have implications for the therapeutic approaches to tackle vascular damage in severe COVID19 disease.

## Materials and methods

### Whole-transcriptome analysis with total RNA sequencing (RNA-Seq)

RNA-seq data files with gene quantifications for primary human epithelial and endothelial cells were downloaded from the ENCODE database (www.encodeproject.org)^6^, using Bioconductor package “ENCODEExplorer” and R. For consistency, only total RNAseq data generated by the same source, “Thomas Gingeras, CSHL”, and aligned to GrCh38 reference genome were included in the analysis. Cell types with one replicate were excluded. Raw counts were converted to counts per million (CPM), filtered based on expression, log2 transformed and normalized with Trimmed Mean of M-value (TMM) method using Bioconductor package “EdgeR”. Only genes with more than 0.3 counts per million (CPM) in at least three samples were kept for the analysis (19360 genes).

The accession numbers for the ENCODE files are ENCFF592BLV, ENCFF788JHJ, ENCFF235TGN, ENCFF401NRN, ENCFF207VFL, ENCFF623ZJJ, ENCFF620THF, ENCFF699QEJ, ENCFF797YZO, ENCFF985PDI, ENCFF110UGQ, ENCFF764AOQ, ENCFF325DPM, ENCFF564EGR, ENCFF233LYV, ENCFF378BDR, ENCFF555QVG, ENCFF972UYG, ENCFF511TST, ENCFF711SNV, ENCFF037WEH, ENCFF498AYE, ENCFF577VBY, ENCFF604VVJ, ENCFF177SUW, ENCFF456RET, ENCFF145DRX, ENCFF224ZRP, ENCFF176IZX, ENCFF674SRB, ENCFF580CHD, ENCFF709VCC, ENCFF747XTG, ENCFF756RKP, ENCFF060LPA, ENCFF262OBL, ENCFF091SWU, ENCFF224YSC, ENCFF819IDA, ENCFF894GLT, ENCFF231GYQ, ENCFF836KPM, ENCFF361WEZ, ENCFF592KDP.

### Endothelial cell cytokine treatments and real-time quantitative polymerase chain reaction

Pooled human umbilical vein endothelial cells (HUVEC; Lonza C2519A) were cultured in Endothelial Cell Growth Medium-2 media (EGM-2) (Lonza). HUVEC were plated at a density of 10,000 cells/96-well plate in EGM-2 supplemented with recombinant human TNF-α (300-01A, PeproTech EC Ltd), IL1-β (200-01B, PeproTech EC Ltd), IL8 (72aa, monocytes derived, 200-08M, PeproTech EC Ltd) and IL-6/IL-6R Alpha Protein Chimera (8954-SR, R&D Biotechne) at 0, 0.01, 0.1 or 1.0 ng/ml for 4 or 24 hours. Total RNA was isolated by using the RNeasy kit (Qiagen) and reverse transcribed into cDNA using Superscript III Reverse Transcriptase (Invitrogen). Quantitative real-time PCR was performed using PerfeCTa SYBR Green Fastmix (Quanta Biosciences) on a Bio-Rad CFX96 system. Gene expression values were normalized to GAPDH expression. Sequences of the human primers used in this study are listed below: *GAPDH* forward: 5’-CAAGGTCATCCATGACAACTTTG-3’ and *GAPDH* reverse: 5’-GGGCCATCCACAGTCTTCTG-3’; *ACE2* forward: 5’-ACAGTCCACACTTGCCCAAAT-3’ and *ACE2* reverse: 5’-TGAGAGCACTGAAGACCCATT-3’.

### Human heart single cell RNAseq data analysis

Human heart single cell raw counts data and the cell annotation meta data were obtained from a recent published study^2^. The data were processed using Seurat package (version: 3.1.1) in R. The endothelial cells which contain *ACE2* reads were grouped as *ACE2*^+^ EC cells, and the rest as *ACE2*^-^ EC cells. The genes enriched in *ACE2*^+^ EC cells were identified using FindMarkers function in Seurat. It applies a Wilcoxon Rank Sum test and then performs Bonferroni-correction using all genes in the dataset. The corrected p value < 0.05 was used as cutoff for significance. To visualize the top 50 genes among different cell groups, the DotPlot function was applied. The size of the dot represents percentage of cells expressing the gene and color represents the scaled average expression levels.

### Epigenetic analysis of chromatin state

Genome-wide ChIP-seq data for histone modifications and DNase I hypersensitivity in HUVEC were obtained from the ENCODE^6^/Broad Institute under GEO accession number GSE29611. ChIP-seq reads were mapped to the human reference genome GRCh37/hg19. Tracks were visualized using the UCSC Genome Browser database (https://genome.ucsc.edu). HUVEC chromatin-state discovery and genome annotation was obtained using ChromHMM^7^ from ENCODE.

### Viral replication assays

#### HCoV-229E

Stocks of HCoV-229E GFP reporter virus described by Cervantes-Barragan et al^4^ were generated following cultivation on the HUH7 cell line. HPAEC (Lonza) and HCAEC (Promocell) were seeded in 96 well plates (2.1×10^4^ cells/cm^2^) 24 hours prior to infection with HCoV-229E-GFP at MOI = 0.6. Virus inoculum was then replaced 1 hour later with 100ul fresh media before incubation at 34°C/5%CO2. GFP fluorescence was measured every 2 hours from 20 to 58 hours post inoculation using the ClarioStar plate reader (BMG).

#### SARS-CoV-2

A sample from a confirmed COVID-19 patient was collected by a trained healthcare professional using combined nose-and-throat swabbing. The sample was stored in virus transport medium prior to cultivation and isolation on Vero E6 (ATCC CRL-1586) cells. Samples were obtained anonymised by coding, compliant with Tissue Governance for the South East Scotland Scottish 279 Academic Health Sciences Collaboration Human Annotated BioResource (reference no. SR1452). Virus sequence was confirmed by Nanopore sequencing according to the ARCTIC network protocol (https://artic.network/ncov-2019), amplicon set V3, and validated against the patient isolate sequence. For the virus isolate used in this project is EDB-2 (Scotland/EDB1827/2020, UK lineage 109, B 1.5) genetic stability was observed up to 5 passages on Vero E6 cells, with particular attention to the S1/S2 furin cleavage site. HCAEC (Promocell) and HPAEC (Lonza) were seeded in a 12-well plate (2.0×10^4^ cells/cm^2^) one day prior to infection with SARS-CoV-2 EDB-2, passage 2, at MOI = 100 or 10 (assessed by endpoint titration on Vero E6 cells). At 0, 24 and 48 hours post infection (hpi) 10 μl of supernatant was lysed in VL buffer (20mM Tris-HCl pH 7.5, 300mM NaCl, 2.5% IGEPAL CA-630, 1:200 RNAsin Plus). The lysate was analysed by RT-qPCR, using GoTaq 1-Step RT-qPCR (Promega) with the SARS-CoV-2 CDC N3 primers (F – GGGAGCCTTGAATACACCAAAA, R – TGTAGCACGATTGCAGCATTG) at 350nM each to determine viral copy number. To assess viral copy numbers, resulting Cts were analysed against a standard curve of lysate with known TCID50/ml value as well as an RNA template.

## Acknowledgements and funding

This research was supported by grants from the Imperial College COVID response fund, the Imperial College British Heart Foundation (BHF) Research Excellence Award and the NIHR Imperial Biomedical Research Centre (BRC) (AMR; GB; MN); BHF/DZHK grant (SP/19/1/34461) and Chan Zuckerberg Initiative (grant number 2019-202666) (MN); the Swedish Science Council, The Cancer Society, the Knut and Alice Wallenberg and Erling Persson Foundation (CB, UL); the University of Edinburgh BHF Research Excellence Award; BHF Chair of Translational Cardiovascular Sciences and H2020 EU grant COVIRNA (Agreement DLV-101016072); IMI-2 CARE (CTB; JGH); BBSRC Institute Strategic Programme grant funding to The Roslin Institute (BBS/E/D/20241866, BBS/E/D/20002172, and BBS/E/D/20002174) (CTB).

## Footnotes

The data, analytic methods, and study materials will be maintained by the corresponding author and made available to other researchers on reasonable request.

